# Improving the accuracy of bulk fitness assays by correcting barcode processing biases

**DOI:** 10.1101/2023.10.31.565047

**Authors:** Ryan Seamus McGee, Grant Kinsler, Dmitri Petrov, Mikhail Tikhonov

**Affiliations:** Department of Physics, Washington University, St. Louis, MO 63130; Department of Bioengineering, University of Pennsylvania, Philadelphia, PA 19104; Department of Biology, Stanford University, Palo Alto, CA 94305

**Keywords:** fitness, sequence barcodes, high-throughput assays, systematic bias, inference

## Abstract

Measuring the fitnesses of genetic variants is a fundamental objective in evolutionary biology. A standard approach for measuring microbial fitnesses in bulk involves labeling a library of genetic variants with unique sequence barcodes, competing the labeled strains in batch culture, and using deep sequencing to track changes in the barcode abundances over time. However, idiosyncratic properties of barcodes (e.g., GC content) can induce non-uniform amplification or uneven sequencing coverage that cause some barcodes to be over-or under-represented in samples. This systematic bias can result in erroneous read count trajectories and misestimates of fitness. Here we develop a computational method for inferring the effects of processing bias by leveraging the structure of systematic deviations in the data. We illustrate this approach by applying it to fitness assay data collected for a large library of yeast variants, and show that this method estimates and corrects for bias more accurately than standard proxies, such as GC-based corrections. Our method mitigates bias and improves fitness estimates in high-throughput assays without introducing additional complexity to the experimental protocols, with potential value in a range of experimental evolution and mutation screening contexts.

## Introduction

Standard assays for measuring microbial fitness involve tracking the abundances of variants over time in batch culture competitions and using this data to estimate relative growth rates (1). High-throughput sequencing technology makes it possible to measure the fitnesses of many strains simultaneously using batch culture assays (2, 3, 4). In this approach, a collection of variants are labeled with unique sequence barcodes for identification (Figure 1A). The pooled variant library is then competed against a reference strain (e.g., the ancestor) over multiple growth cycles (Figure 1B). The batch culture is sampled at designated intervals, and barcode regions are extracted, amplified, and sequenced for each sample. The relative abundance of a variant at a given time can be determined from the fraction of the total sequencing reads that map to its barcode in the corresponding sample. In the simplest approach, the relative fitness of each variant can be estimated by fitting a linear model to its log-count trajectory (Figure 1C).

**Fig. 1.**
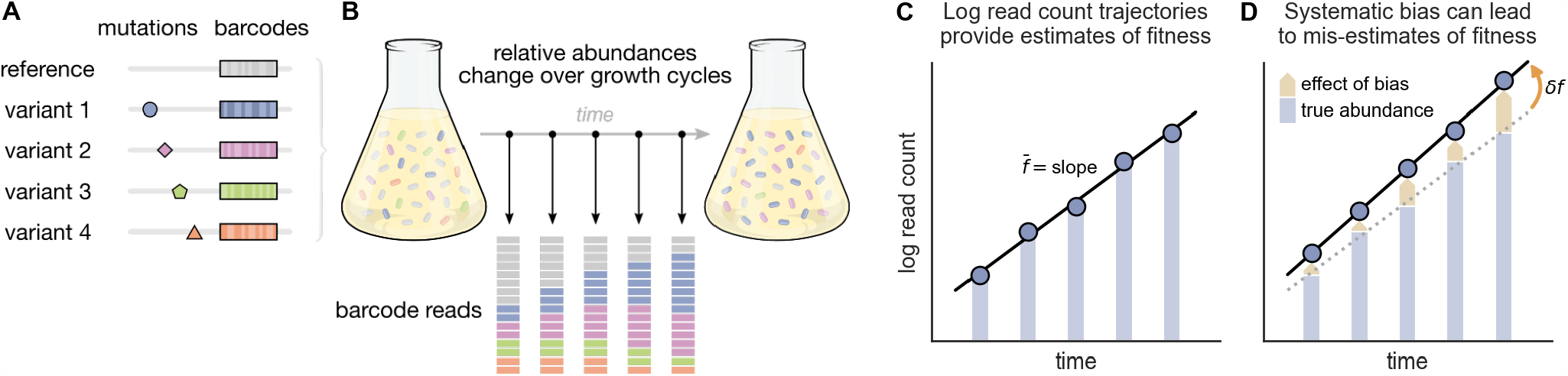
Genetic barcodes that are susceptible to systematic amplification and sequencing biases can cause misestimates of fitness. (**A**) A standard approach for bulk fitness assays involves labeling a library of genetic variants with unique sequence barcodes. (**B**) The variant library is pooled and grcwn in batch culture, often over multiple serial dilution growth cycles. Samples of the batch culture are taken at designated time points, and barcode sequences are extracted, amplified, and sequenced for each sample. The frequency of a barcode among all sequencing reads in a sample provides an estimate of the corresponding variant’s relative abundance in the batch culture at that time. (**C**) The relative abundances of variants are expected to change exponentially over time according to their relative fitnesses. As such, the log read count of each variant’s barcode is expected to change linearly, where the slope of the best-fit line presides an estimate of the variant’s fitness 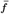 (l.e., exponential growth rate). (**D**) However, bias-inducing factors in the amplification and sequencing process may cause barcode counts to be under-or ever-represented relative to the true abundance of the corresponding variants in culture, which can lead to a misestimation of fitness, *δf*.

This approach provides good estimates of fitness when barcode read counts are a reliable reflection of the relative abundances of variants in the culture. However, variability in read counts can be introduced by multiple factors. Some noise in read counts is to be expected due to stochasticity in growth cycles, perturbations in culture conditions (e.g., ‘batch effects’ such as incubation temperature, nutrient concentrations, or inoculum densities), and bottlenecks associated with serial transfer and sampling. However, uncertainty in fitness estimates attributable to random noise can typically be quantified and mitigated through suitable control strategies.

Here we are concerned with sources of bias that cause barcode read counts to systematically deviate from the variants’ true culture abundances. While genetic barcodes are often assumed to be inert labels, it has been shown that their sequence properties can lead to non-uniform amplification and uneven coverage in next-generation sequencing pipelines (5, 6, 7). For example, the base composition of a barcode can modulate its sensitivity to small fluctuations in temperature ramps and enzyme activities during PCR amplification, which can cause some barcodes to be consistently over-or under-represented in samples (8, 9). This can introduce systematic biases that result in erroneous read counts and misestimates of fitness (Figure 1D). Barcode representation bias is known to correlate with statistics such as GC ratio (8, 9, 10, 11, 12, 13), but the relationship between amplicon sequence and bias is more complex than GC content alone and has not been fully characterized (5, 14). Furthermore, amplification bias also depends on the idiosyncratic processing conditions for each sample, which makes it challenging to determine the extent to which counts and fitness estimates have been impacted by bias (15).

One strategy to mitigate barcode-associated bias is to label each variant with multiple distinct barcodes such that their differing biases average out when taken as an ensemble. However, the addition of redundant barcodes can introduce substantial complexity to library preparation, which limits the scalability of this approach. Beyond this, it is not straightforward to label variants with redundant barcodes in many experimental evolution contexts, such as in lineage tracking experiments where a clonal ancestral population is prelabeled with random barcodes before *de novo* mutants are spontaneously generated (16, 17).

Here we present a data-driven method that infers and removes the effects of barcode processing bias by leveraging the structure of systematic error across an experimental data set. This method offers a procedure for correcting bias and improving fitness estimates from high-throughput assays without introducing complexity to libraries or assay protocols.

## A Data-driven Approach

We aim to infer and correct systematic biases using the data that is typically collected in bulk fitness assays, namely barcode read counts for a library of variants measured across time series of competition culture samples. Often the same barcode-labeled library is assayed multiple times, either as biological replicates or to measure fitnesses in alternative environmental conditions. The involvement of the same barcodes in multiple assays enhances the accuracy of our bias inference, but our method can also be applied to single-assay data.

For narrative simplicity, we assume that sequencing depth (i.e., total read count) is the same for all samples, and we refer simply to barcode counts rather than “normalized counts” or “relative abundances” (Supplementary Section S1.1). Sequencing depth is never actually uniform, but correcting for varying sequencing depth is standard practice (in fact, our algorithm has a built-in capacity to do so; see Supplementary Section S2.1b). We further assume that the variants of interest make up a small fraction of the batch culture relative to the reference strain, such that their change in abundance during an assay is well-modeled by an exponential. Making this assumption and accounting for deviations from it are also standard practice. Under these assumptions, the log-counts of each barcode are expected to follow linear trajectories, which simplifies the presentation of our method.

### Model of underlying bias

In practice, observed counts reflect not only the relative abundances of variants, but also the effects of noise and bias that arise in the barcode amplification and sequencing process. Here we are concerned with the contributions of barcode processing bias in particular. This bias differs from random noise in that its effects on observed counts will tend to impact a given barcode in a systematic way across samples. That is, a variant whose barcode is highly susceptible to bias will tend to have its counts affected more strongly across all samples, compared to other variants. Similarly, samples with procedural conditions that induce substantial bias will exhibit greater impacts on variants’ counts across the board when compared to other samples.

We model the effect of bias 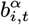 on the observed count of variant *i* at time *t* in assay *α* as the product of the variant’s characteristic susceptibility to bias *u*_*i*_ and the prevalence of bias in that sample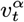

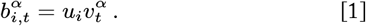

The bias prevalence component gives the overall strength and direction of bias in a sample, which reflects the tendency of the procedural conditions in that sample to influence barcode counts. In this model, the susceptibility of a variant’s barcode modulates how much that variant’s counts are affected by processing bias. Prevalent bias translates to relatively large shifts in counts for highly susceptible variants (Figure 2A, top and bottom variants), while variants with negligible susceptibility are weakly affected by bias-inducing factors, even when they are highly prevalent (Figure 2A, middle variant). The counts of ‘positively susceptible’ barcodes are influenced in the opposite direction from ‘negatively susceptible’ barcodes with respect to over-versus under-representation (Figure 2A, compare top and bottom variants).

**Fig. 2.**
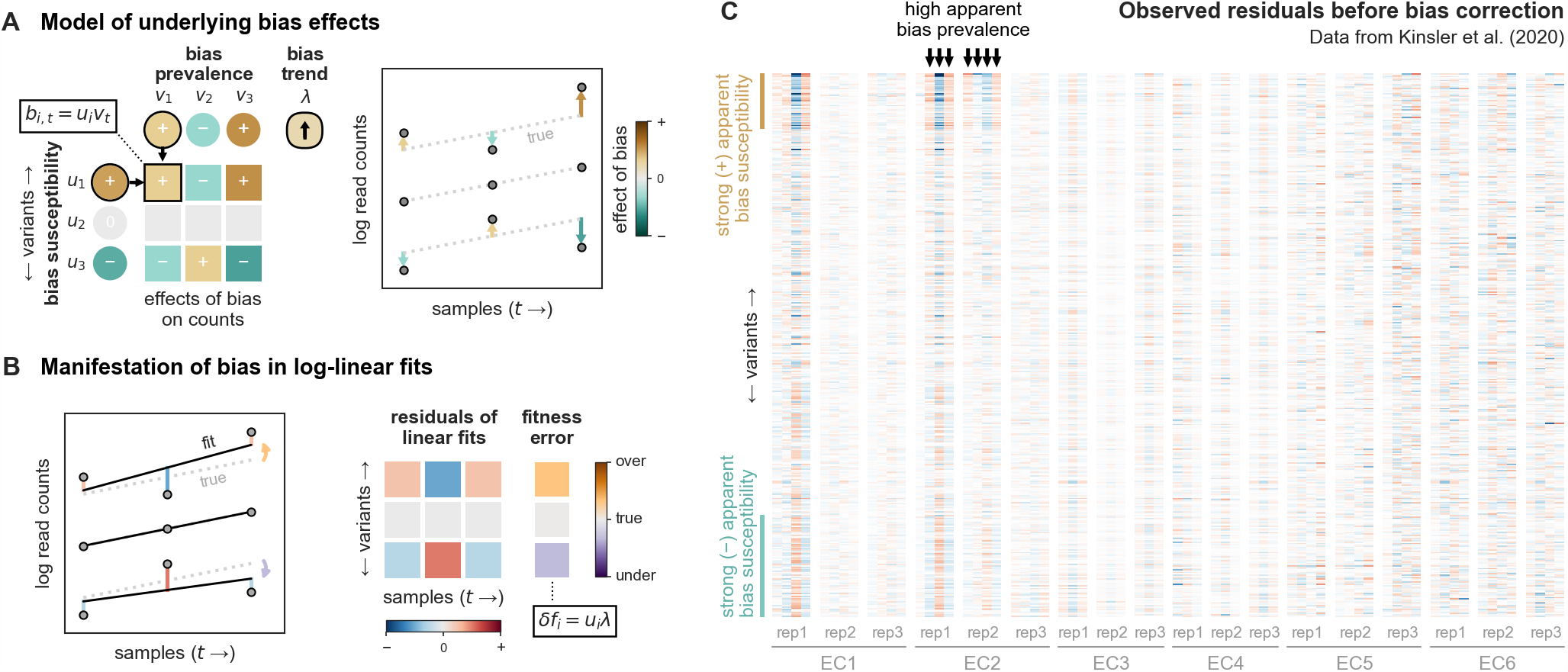
Barcode processing bias Impacts the structure of residuals and the accuracy of fitness estimates derived from linear fits of log-count trajectories. (**A**) Log-count trajectories are shown for a hypothetical fitness assay with three variants (right). The contributions of barcode processing bias to the observed counts are shown in the table (left). We model the effect of bias on the count for variant *i* in sample *t* as the product of the prevalence of bias in the sample (*v*_*t*_ values, one per sample) and the variant’s susceptibility to bias (*u*_*i*_ values, one per variant). These bias effects cause observed counts (points, right) to deviate (arrows, right) from the log-linear trajectories that correspond to each variant’s true change in abundance over time (dotted lines, right). (**B**) A line Is fit to each variant’s observed log-count trajectory (black lines, left). The residuals of each fit are depicted as bars (left) and in the table (right). The effects of underlying bias are reflected in the structure of the magnitudes and signs of residuals (refer to the color scale below the table). Note the similarities between the bias effect table in (**A**) and the residuals table in (**B**). (**C**) The heatmap shows the residuals of linear fits to log-count trajectories for a library of mutants (rows) across 18 fitness assays (major column groups labeled as replicates) conducted in six different “evolutionary conditions” (EC) batches from Kinsler et al. (18)(see Supplementary Section S3 for more information); color scale same as in (B). Variants (rows) are ordered by the GC ratio of their barcodes, decreasing from top to bottom. We observe that barcodes (rows) with extreme GC ratios tend to have large residuals across multiple samples, which is consistent with these variants being highly susceptible to bias (example blocks of variants indicated with brown and teal bands to left of table). Samples (individual columns) with relatively large residuals and a strong correlation between residuals and GC ratios are consistent with high bias prevalence (example samples indicated by arrows above table).

### Manifestation of bias in misestimates of fitness

The log-count of each variant’s barcode is expected to change linearly with a slope that corresponds to its growth rate. Fitting a line to the log-count trajectory of each barcode provides an estimate of the relative fitness 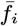 for each variant (Figure 1C). However, barcode processing biases cause read counts to deviate from the “true” trajectories. Under our model of bias, we expect these effects to leave two notable signatures in the data.

First, bias often perturbs observed counts away from tightly log-linear trajectories. As a result, the residuals of a variant’s log-linear fit often reflect the magnitudes and directions of bias effects in the respective samples (note the correspondence between the underlying bias effects in Figure 2A with and the observed residuals in Figure 2B). Barcodes that are highly susceptible to bias are expected to have large residuals in samples where bias is prevalent (Figure 2B, top and bottom variants), whereas barcodes that are not susceptible to bias will be immune from these deviations (Figure 2B, middle variant).

Patterns of bias-driven residuals can be seen in the experimental data set from Kinsler et al. (18) shown in Figure 2C. This study collected barcode read count time series for more than 500 variants across a number of bulk fitness assays (a subset of which are shown in Figure 2C, see Supplementary Section S3 for more information). We see that some of these samples (e.g., columns marked by arrows in Figure 2C) exhibit strong residuals, where the sign and magnitude of residuals also appear to be correlated with the GC ratio (i.e., row ordering) of the respective barcodes. This structure is consistent with systematic barcode processing bias being prevalent in these samples, in contrast to other samples where residuals are more random across variants. In those samples where systematic bias appears to be prevalent, barcodes with extreme GC ratios (e.g., brown and teal bands in Figure 2C) tend to have larger residuals, suggesting that these variants are more susceptible to the effects of this bias. However, the GC ratio of a barcode does not always correspond to the apparent susceptibility (i.e., residual magnitude) of variants. This is expected because barcode processing bias is related to a number of sequence properties, with GC content being just one correlate.

The second major effect that bias-induced shifts in counts can have is to cause the best-fit slope of a variant’s log-count trajectory to deviate from its actual rate of change in abundance (Figure 2B). This occurs when the effects of bias vary from one sample to the next such that a variant’s barcode becomes increasingly over-or under-represented over the course of an assay (such as in Figure 1D). Such a trend in bias confounds the signal from a variant’s change in abundance and results in a misestimate of fitness (note that bias with a constant effect on all counts in a time series shifts the log-linear fit vertically but does not change its slope). The degree to which a variant’s fitness estimate is impacted by a trend in bias prevalence is modulated by its bias susceptibility (Figure 2B).

We can formalize the temporal trend of bias prevalence in an assay with a linear model

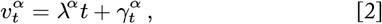

where λ^*α*^ gives the change in prevalence over time and 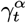 is the incidental deviation from this linear trend for each individual sample. Then, the error in a variant’s fitness estimate relative to its true fitness 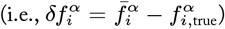 is given by the product of the variant’s bias susceptibility and the trend in bias prevalence in that assay (Supplementary Section S1.3):

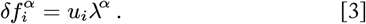

Unfortunately, it is impossible to disambiguate bias-driven trends in counts from fitness-driven ones using counts or residuals alone. Nevertheless, as we will explain in the next section, having just *one* control group of variants with equal true fitnesses is sufficient to fully resolve this problem and infer the effects of bias for an entire library.

### Disentangling bias trends from fitness estimates

Consider the thought experiment presented in Figure 3, where the ground truth fitnesses, bias susceptibilities, and sample bias prevalences underlying a three-assay data set are known. Here, we consider a set of neutral variants that all have the same fitness as the reference strain (e.g., the ancestor) and whose relative abundances thus remain constant throughout each assay. Nevertheless, we see fluctuations in barcode read counts (grayscale tables in Figure 3A) due to the effect of bias in each sample.

**Fig. 3.**
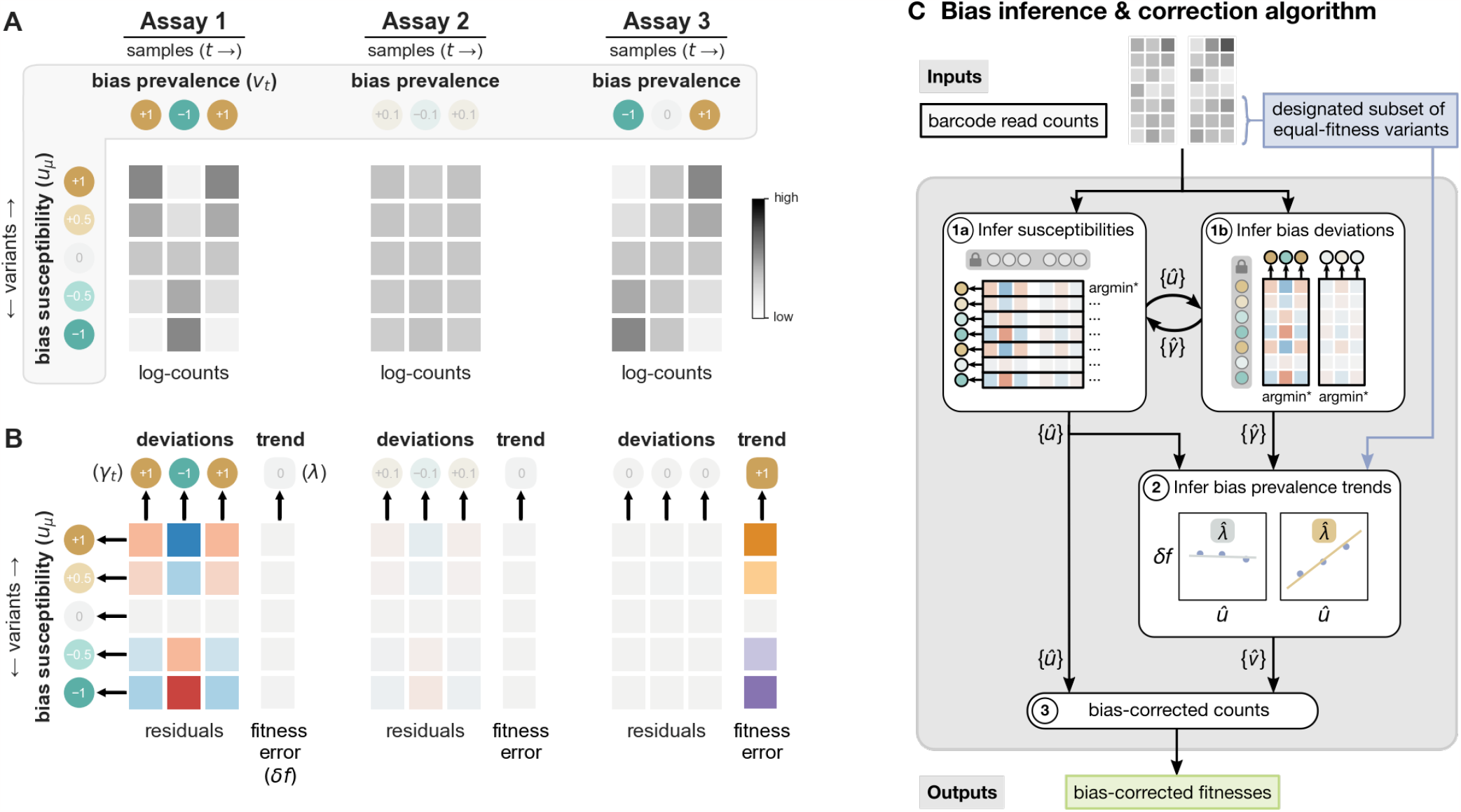
Decomposing bias prevalence components into trends and deviations enables their inference. (**A**) Log-counts from three hypothetical assays involving a set of five variants are shown (gray-scale tables). For illustrative purposes, these five variants and the reference strain are assumed to all have the same fitness. Therefore, the abundances of these variants in culture remain constant over time. However, observed counts are seen to deviate from constant trajectories due to the effects of barcode processing bias. The ground truth bias susceptibility and bias prevalence components that give rise to the observed counts are shown in the gray outlined region. (**B**) A line is fit to each observed log-count trajectory, and the residuals of each fit are depicted in the tables for each assay (below). The slope of each linear fit provides a fitness estimate for the corresponding variant. The errors in fitness estimates due to bias-induced shifts in log-count trajectories are given in the columns to the right of each residuals table. These linear fits effectively decompose counts data into trends (fitnesses) and deviations (residuals). Decomposing bias prevalence into trend (λ) and deviation (*γ*_*t*_) components as well enables us to infer these components using the analogous terms derived from the counts data. Black arrows indicate that i) inference of bias susceptibility values is informed by each variant’s set of residuals across samples, ii) inference of bias prevalence deviations is informed by each sample’s set of residuals over all variants, and iii) inference of bias prevalence trends is informed (in part) by fitness misestimates. (**C**) A graphical schematic of the two-stage bias inference algorithm is shown. See the text and Supplementary Section S2 for more information.

In this example, each variant’s pattern of residuals across all assays (i.e., the signs and magnitudes in each row of residuals) tends to correspond to the strength and direction of that variant’s underlying susceptibility. Similarly, underlying bias prevalence values are often closely associated with the patterns of residuals across variants (i.e., columns of residuals).

However, the relationship between bias prevalence and residuals is more subtle, as is evident when comparing the three assays in this example. In Assay 2, bias prevalence is low in all samples, all residuals are correspondingly small, and fitnesses are accurately estimated. In Assay 1, bias prevalence is greater, and susceptible variants are more impacted as seen in larger residuals for those variants. However, fitness estimates remain accurate because the bias does not have a temporal trend that confounds variant growth over time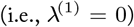. Compare these outcomes with those of Assay3, where bias prevalence increases precisely linearly over time 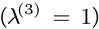. This results in log-linear count trajectories that are perfectly fit by lines with zero residuals (in the absence of other noise), but the apparent slopes are solely attributable to the the trend in bias and therefore give erroneous fitness estimates. This illustrates why residuals alone are not sufficient to infer bias prevalence without additional information about the trend in bias for each assay.

Notice that we have decomposed both log-counts and bias prevalence values into trend (slope) and deviation (residual) terms (Figure 3B). Sets of residuals across variants in a sample are informative about the respective *deviation* of that sample’s bias prevalence from the overall trend in prevalence for the assay (note the correspondence between columns of residuals and bias prevalence deviation values, 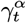, for all assays in Figure 3). It follows that fitness estimates provide information about *trends* in bias prevalence.

Equation 3 tells us that we can quantify the trend in bias prevalence *λ*^*α*^ by regressing fitness errors 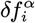 against bias susceptibilities *u*_*i*_. In general, it is circular to assume that one would know by how much fitnesses have been misestimated, since the purpose of the assay is to measure fitnesses in the first place. However, if a subset of variants are known beforehand to have the same fitness (e.g., a set of genetically identical strains labeled with different barcodes), then fitness errors can be approximated for this subset by comparing the apparent fitness of each variant to the mean estimate of the group. Therefore, a *single subset* of equal-fitness variants provides the information necessary to estimate the trend in bias prevalence that is experienced by *all* variants in the library.

Altogether, the logic of our method for inferring bias terms is as follows: i) the magnitudes and signs of a variant’s collection of residuals across all available samples inform the magnitude and sign of that variant’s bias susceptibility; ii) the magnitudes and signs of a sample’s collection of residuals over all variants inform the magnitude and sign of that sample’s deviation in bias prevalence from a general trend in prevalence over the respective assay; and iii) the correlation of fitness misestimation with bias susceptibility among a control subset of equal-fitness variants informs the trend in bias prevalence itself. The inferred bias components are used to compute bias-corrected counts that no longer include spurious trends that confound estimates of fitness.

### The bias inference algorithm

Our method infers underlying bias components from barcode read count time series data in two major stages.

First, we infer bias susceptibility values and bias prevalence deviation values using an iterative optimization process. A particular set of bias susceptibility and bias prevalence estimates(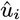 and 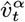, respectively) generates a corresponding set of bias-adjusted counts

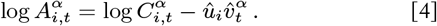

Each adjusted count time series can be fit by a log-linear model

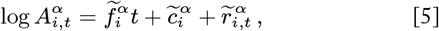

Where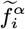 gives the adjusted fitness estimate for variant *i* in assay *α*,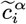 gives the y-intercept, and 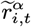 denotes the residual of each sample *t* from the log-linear fit (Supplementary Section S1.2). We employ the heuristic that accurate estimates of bias susceptibility and prevalence deviations will yield bias-corrected counts that minimize residuals across the data.

We begin by initializing bias susceptibility and prevalence terms to random values (Supplementary Section S2.0). To infer the bias susceptibility of variant *i*, we fix the set of bias prevalences at their current values and solve for the susceptibility value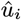 that minimizes the set of residuals associated with that variant across all assays and samples in the data set (Figure 3C, box 1a). Evaluating this optimization for every variant in turn produces an updated set of bias susceptibility values for the library. Next, we fix the set of variant susceptibilities at their current values and infer the bias prevalence deviation terms 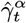 that minimize the set of residuals in each assay *α* across all variants (optimization of bias prevalence deviation values is done on a per-assay basis; Figure 3C, box 1b). We alternate between optimizing bias susceptibility values (given the current prevalence deviation values) and optimizing bias prevalence deviation values (given the current susceptibility values) until suitable convergence is reached (regularized objective functions are used to resolve symmetries and ensure convergence, see Supplementary Section S2.1).

In the second stage of our process, we infer the trend in bias prevalence for each assay in order to determine absolute estimates of bias prevalence. Following the logic derived from Equation 3, we estimate the trend in bias prevalence in an assay *α* by regressing fitness misestimates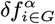 against inferred bias susceptibilities 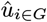 for a group of variants *G* where these values are known (Figure 3C, box 2). In particular, we perform this step using a designated subset of variants that are known beforehand to have the same fitness, which allows us to estimate their fitness errors relative to the group mean. The slope of this regression provides an estimate of the trend in bias prevalence for the respective assay (see Supplementary Figure S6 for examples). With inferred values for both the trends and deviations in bias prevalence in hand, we then compute absolute bias prevalence estimates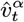 for each sample (Supplementary Section S2.2).

The inferred bias susceptibility and bias prevalence values we obtain from this procedure allow us to estimate and correct the effect of bias for each individual count in the data set (Equation 4). More information about the implementation of this algorithm is provided in Supplementary Section S2.

## Results

We applied our bias inference and correction method to the bulk fitness assay data from Kinsler et al. (18) that was introduced above (Figure 2C). The data used in this analysis include barcode read counts for a library of 549 yeast variants across 35 assays that feature a range of culture conditions (see Supplementary Section S3 for more information).

This variant library is derived from a collection of adaptive yeast mutants that were generated in a prior evolution experiment (16, 17). This library includes a large diversity of mutant genotypes, but several subsets of variants with similar mutations and corresponding fitness effects can be identified (Supplementary Section S3.2). For example, many mutants underwent a whole-genome duplication to become diploid but have otherwise very similar genetic backgrounds. The fitness effect of genome duplication is similar for all diploid mutants across the assays considered here (Supplementary Section S3.2). Similarly, two other subsets of variants have mutations in the same gene—*GPB2* and *PDE2*, respectively—that confer similar fitness effects across assays (Supplementary Section S3.2). Here we consider these variant subsets to be phenotypically identical and treat them as equal-fitness reference groups in our analysis. We use the Diploid group as the equal-fitness control group for inferring temporal trends in bias prevalence with our method. The *GPB2* and *PDE2* groups serve as independent validation groups for assessing the degree to which our method improves the accuracy of fitness estimates.

Results of bias inference using our method on this data set are shown in Figure 4A (shown are 18 representative assays across six “evolutionary conditions” (EC); results for 17 additional assays can be found in Supplementary Section S3.4). The inferred bias prevalences are consistent with expectations regarding which samples exhibit systematic bias based on the patterns of residuals observed in the original data (Figure 2C). Bias-corrected counts yield improved linear fits, as evident in the overall decrease in the magnitude of residuals after correction (Figure 4A; compare to the pre-correction residuals in Figure 2C and Supplementary Figure S7). Our method’s corrections effectively reduce the structural patterns of residuals across samples and variants, which is consistent with mitigating the effects of systematic bias.

**Fig. 4.**
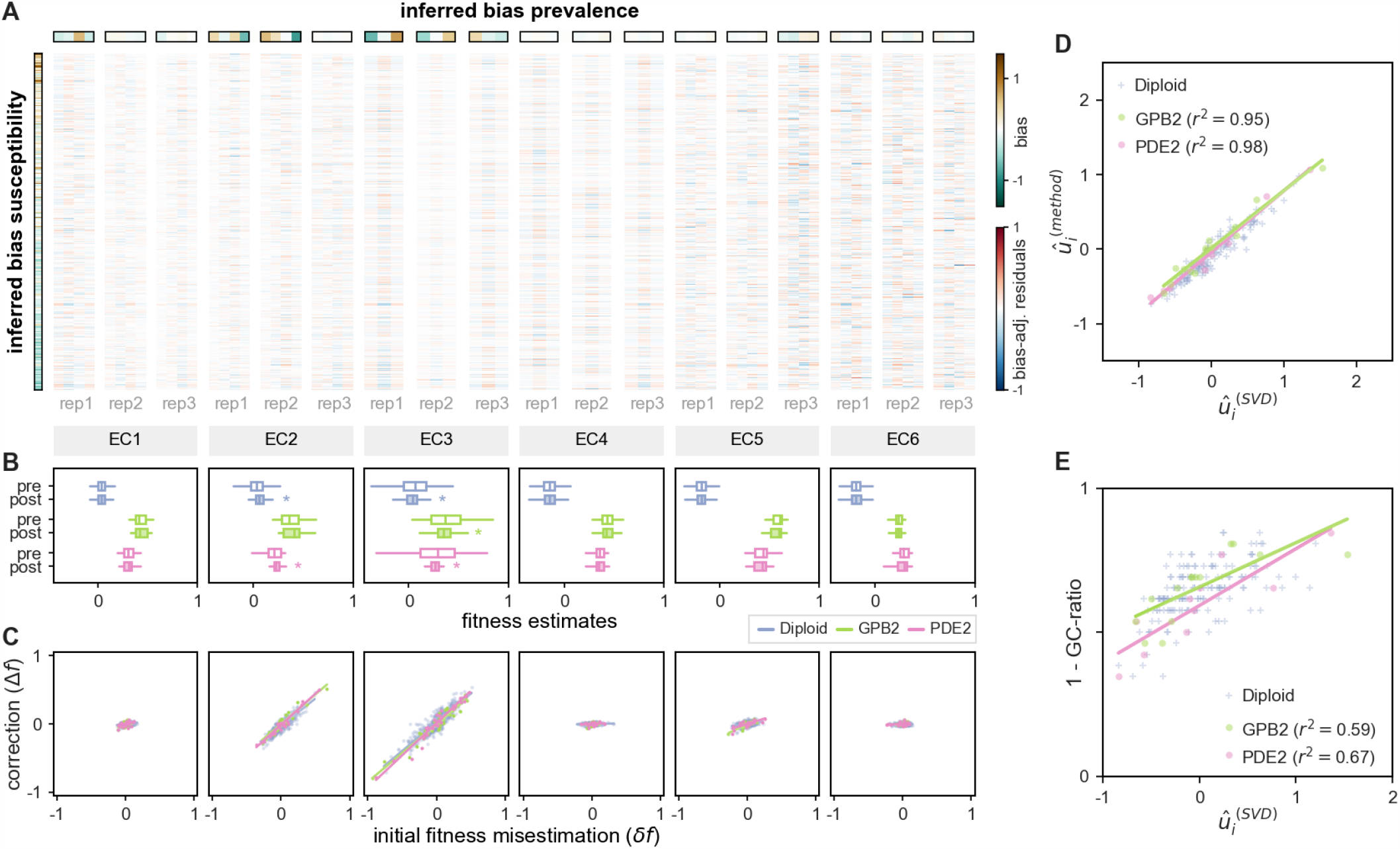
Our method infers bias terms that enable bias reduction and more accurate fitness estimates. We applied our method to the counts data from Kinsler **et al**. (18) that was introduced in Figure 2C. (**A**) The bias susceptibility and bias prevalence values inferred by our method are shown in the outlined bands to the left of and above the table, respectively. The post-correction residuals of log-linear fits to the corresponding bias-corrected counts are shown in the table. These bias-corrected residuals have reduced magnitude and structure compared to the pre-correction data shown in Figure 2C. (**B**) The pre- and post-correction distributions of fitness estimates (white and shaded boxplots, respectively) are shown for each reference group of variants across six growth conditions. The variance of fitness estimates decreases significantly (statistical significance, denoted by *, is determined by Levene’s test for equal variances; *p* < 0.05) in the EC2 and EC3 conditions, for which multiple assays have high bias prevalence with strong temporal trends. (**C**) The correspondence between each variant’s initial fitness misestimation using counts before bias correction 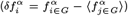 and the change in its fitness estimate following bias-correction 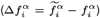 is depicted using scatter plots for each growth condition (each point represents a variant). (**D**) The bias susceptibility values returned by our method are highly correlated with bias susceptibilities inferred using an independent, SVD-based method, which we consider to be approximate ground truth values. (**E**) The approximate ground truth bias susceptility of each variant is correlated with the GC ratio of its barcode, as expected, but this correlation is weaker than it is for the bias susceptibility values returned by our method (seen in (D)).

The impacts of our method’s bias corrections on fitness estimates for the Diploid, *GPB2*, and *PDE2* reference groups are shown in Figure 4B. Variants in each group are expected to have the same fitness, so within-group variation in pre-correction fitness estimates is likely attributable to barcode processing bias. Our method significantly reduces the variance of fitness estimates in assay conditions where within-group variation is relatively high before correcting for bias (e.g., EC2, EC3). We see that significant reductions in fitness estimate variance coincide with assays where bias is both relatively prevalent and has a strong temporal trend (e.g., bias prevalence going from strongly negative to strongly positive, or vice versa). In conditions where fitness estimates are already tight in the original data, our method tends to infer low bias prevalence and does not significantly alter either the mean or variance of fitness estimates.

Figure 4C provides a closer look at how our method’s bias corrections impact fitness estimates at the level of individual variants. In conditions where initial fitness estimates have substantial errors (e.g., EC2, EC3), the updates to fitness estimates induced by our bias-corrections 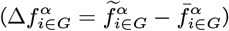 are highly correlated with the initial errors in estimation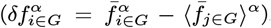.This demonstrates that the corrections made by our method address the underlying cause of misestimation for each particular variant. On the other hand, our method does not significantly alter fitness estimates in conditions where fitness errors are already low. Therefore, our method successfully corrects fitness misestimation when bias is prevalent and confounding, yet conserves accurate fitness estimates when bias is negligible.

We can further validate our inference by comparing the bias components returned by our algorithm to those determined using an independent method. The relationship given by Equation 3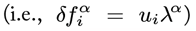 suggests such an alternative. If we collect the fitness misestimates for all variants and all assays in a matrix (rows and columns, respectively), then the structure of this matrix will encode information about the bias susceptibility and prevalence trends that are consistent with the observed errors across the data set (Supplementary Figure S4). Performing singular value decomposition (SVD) on this matrix generates an independent estimate of precisely the bias mode that our method aims to infer. In particular, the first left singular vector provides estimates of bias susceptibility for each variant, and the first right singular vector gives estimates of the trend in bias prevalence in each assay.

We stress that this SVD approach, which we use here for validation, computes bias terms by decomposing fitness *error* data, which requires knowledge of true fitnesses for the collection of variants and assays in question. True fitness information is generally not available beforehand in the context of bulk fitness assays, which prohibits the use of the SVD method for library-wide bias inference. However, the SVD approach can be applied individually to known subsets of equal-fitness variants, such as our Diploid, *GPB2*, and *PDE2* reference groups, for which true fitnesses can be approximated using the subset’s average fitness estimate.

Figure 4D shows a tight correspondence between the variant bias susceptibility values inferred using our method and the corresponding values estimated using the SVD approach. The strong correlation seen in every reference group validates the bias inference performed by our algorithm. Critically, note that our method infers bias susceptibility values for the entire variant library, while the SVD approach can only be applied to special variant subsets for which *a priori* fitness information is available. The ability of our method to infer and correct biases for an entire variant libary over multiple assays simultaneously, while requiring only a single subset of variants as a control, is a significant advantage of our method over other methods that have greater control or redundancy demands (e.g., labeling every genetic variant with multiple barcodes).

In Figure 4A, we see that the variants with the strongest inferred susceptibilities (in either direction) are often those with the most extreme GC ratios, as predicted by the structure of residuals in the original data. That said, a barcode’s GC ratio is only weakly correlated with its susceptibility to bias (Figure 4E). While GC content is known to play a role in modulating how a barcode responds to bias-inducing conditions in the amplification and sequencing process, it is not the only factor that influences a barcode’s representation in a sample (5, 14). Our method’s bias susceptibility values accurately capture each barcode’s response to this multi-factorial bias mode and better account for the systematic structure in the data.

## Discussion

We have developed a computational method for inferring and correcting the effects of barcode processing biases in high-throughput fitness assays. Our method infers the bias susceptibility of each barcode as well as the prevalence of bias in each sample. These components are used to estimate the effect of bias on each individual count, from which bias-adjusted effective counts are obtained. These bias-corrected counts are better representations of the actual changes in variant abundances, and thus provide more accurate estimates of fitness.

Here we model the effects of barcode processing bias as a single error mode (i.e., a single coherent pattern of effects), which we find captures most of the deviations in this data (Supplementary Section S3.2). However, multiple factors may jointly contribute to the observed effects of this bias mode. We see that the GC ratio of a barcode is weakly correlated with its susceptibility to bias (Figure 4E), which suggests that GC content explains some, but not all, of the bias in this data. While we do not characterize all of the factors contributing to barcode processing bias, our method successfully estimates the comprehensive effect of this bias mode on each count. In turn, the estimates of bias susceptibilities and bias prevalences returned by our method may help pinpoint the features of barcodes or protocols that are inducing bias. In contexts where there are multiple independent sources of bias, a model with additional bias modes may account for more of these effects, but our methodology can be readily extended to multiple mode inference where necessary (i.e., by repeating the basic inference algorithm for each bias mode).

A notable advantage of our approach over other strategies for mitigating barcode-associated biases is that our method requires only one multiply-labeled control subset to infer and correct the effects of bias for the entire library. By contrast, other approaches, such as the SVD method considered above (see Results), require many or all genetic variants to be redundantly labeled in order to resolve the impact of bias. Instead, by decomposing bias into its constitutive susceptibility and prevalence components, our method can leverage coherent structure in a data set to infer these effects without barcode redundancy, requiring a single control set only to disambiguate temporal trends driven by bias from those driven by fitness.

In general, the accuracy of our method depends in part on the fidelity of the equal-fitness control set. Ideally, this group would consist of a set of differentially labeled but otherwise genetically identical strains to ensure absolute fitness neutrality among the control variants. It is relatively straightforward to incorporate such an ideal control set into bulk fitness assays *a priori* (e.g., label the reference strain with a collection of barcodes). When such an ideal control set is not feasible (such as when analyzing a pre-existing data set), a subset of variants that are found *a posteriori* to be nearly-neutral can be used, as was done here for the Kinsler et al. (18) case study. However, the ability of the method to disambiguate bias trends from fitness effects will tend to diminish as the magnitude of biological (i.e., fitness) differences among the control variants increases. The accuracy of bias inference will tend to improve as the size of the control subset (i.e., the number of distinct barcodes included) increases, but reasonable accuracy can often be achieved with a modest number of orthogonal barcodes.

Similarly, the quality of bias estimates and corrections using this method scales with the size of the data set. In principle, this method can be applied to data from a single fitness assay. However, given that the strength of this method is in leveraging consistent patterns in the data, its inference will be better constrained and more accurate with multiple assays. It does not matter if the assays represent replicates from the same environment or assays performed in different environments so long as the barcodes and processing protocols (i.e., the potential sources of bias) are the same throughout. The degree to which inference is improved by the inclusion of additional assays depends on how prevalent bias is in the respective assays.

A benefit of our method is that it does not make demands on variant library design or fitness assay protocols, so long as at least one equal-fitness control set is included. This also avails our method as a post-processing tool that can improve the accuracy of fitness estimates with low overhead in many contexts. Therefore, this computational approach offers significant value to a wide range of experimental fields where accurate fitness measurements are of interest, such as experimental evolution, lineage tracking, and deep mutational scanning.

## Author Contributions

Conceptualization: MT. Data Curation RSM, GK, MT. Formal Analysis: RSM, MT. Funding: MT. Investigation: RSM, GK, MT. Methodology: RSM, MT. Project Administration: RSM, MT. Resources: GK, DP. Software: RSM, MT. Supervision: DP, MT. Validation: RSM, MT. Visualization: RSM. Writing (Original Draft): RSM. Writing (Review & Editing): RSM, GK, MT.

The authors declare no conflicts of interest.

## Acknowledgments

We thank Olivia Ghosh, Olivia Kosterlitz, Jacob Moran, and the Tikhonov group for valuable discussions.

## Supplementary Information

### S1 Bias Model Details & Notation

#### S1.1 Barcode read counts and systematic barcode processing bias

We consider data sets collected from bulk fitness assays. These data consist of barcode read count time series collected for a library of variants (indexed by *i*) from one or more assays (indexed by *α*; representing e.g., replicates or alternative environmental conditions). The relative count of a barcode among all counts for a given sample is expected to represent the relative abundance of the corresponding variant in the assay culture. We refer to the set of counts supplied to our algorithm as *raw* counts (although these input counts may have been pre-processed for other reasons). The raw read count of barcode (variant) *i* at time point (sample) *t* in assay *α* is denoted by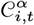.

Several factors can cause raw counts to deviate from relative abundances. First, sequencing depth (i.e., total reads per sample) typically varies from one sample to the next. This can cause sequences of counts to exhibit fluctuations that are not representative of changes in the composition of the population. Therefore, counts must be normalized to account for sequencing depth variation within each assay before time series analyses (e.g., growth rates estimation) can be done. This can be achieved by dividing the counts in each sample (*t*) by a set of per-sample normalization factors 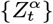 that remove offset differences in total counts across time points:

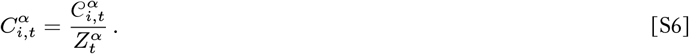

We refer to the resulting normalized counts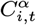 as *observed* counts (distinguishing them from raw input counts and the bias-adjusted counts discussed below). Our method infers appropriate depth-normalization factors as part of the bias inference algorithm (Supplementary Section S2.1b).

Random noise and systematic biases can also cause individual barcode read counts to deviate from accurate representations of relative abundances. Let the “*true*” count of barcode *i* in sample *t* of assay *α* be given by 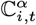 that is, the count value that would be observed in the absence of barcode processing bias or noise. The corresponding count that is actually observed,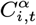, incorporates the “true” barcode representation as well as the contributions of processing bias 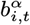and noise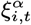 on this data point (focusing on log counts here):

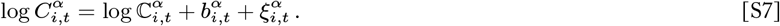

Our model of systematic barcode processing bias posits that the bias can be decomposed into two components: the characteristic susceptibility of barcode *i* to bias, *ui*, and the prevalence of bias-inducing effects in sample *t* of assay 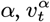 :

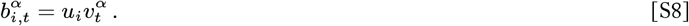

Changes in bias prevalence over the course of an assay can confound estimates of growth rates. To address temporal trends in bias prevalence, we describe the prevalence of bias at time (sample) *t* of assay *α* using a linear model:

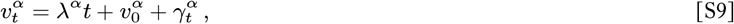

where the slope *λ*^*α*^ gives the overall trend in bias over the course of the assay, and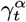 values represent the deviations from this trend for each individual sample (and where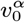 is the y-intercept).

#### S1.2 Linear models of log-counts: Fitness estimates and residuals

We assume that bulk fitness assays are conducted using conditions where the variants of interest segregate exponentially (e.g., variants of interest make up a small fraction of the population relative to the reference strain), or that the raw data has been pre-processed to account for deviations from this assumption. Under this assumption, the log-count of barcode *i* is expected to change linearly with a slope that corresponds to the fitness (growth rate) of the associated variant. Therefore, an estimate of fitness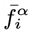 can be obtained by fitting a linear model to the corresponding observed counts:

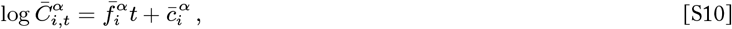

Where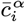 is the y-intercept and log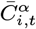 is the predicted log-count of barcode *i* in sample *t* under the best-fit linear model (the ‘bar’ notation denotes quantities inferred from the *observed* counts data). An observed log-count deviates from the corresponding best-fit line by the residual amount 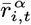:

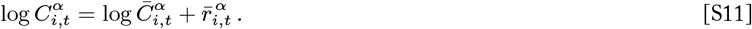

#### S1.3 Mis-estimation of fitness before bias correction

Given that observed counts incorporate the effects of bias and noise, estimates of fitnesses given by the slopes of lines fit to observed count trajectories may deviate from the true fitnesses of variants. Suppose that we had access to “true” counts that were unaffected by bias or noise. In the absence of these deviation effects, true counts would follow perfectly log-linear trajectories with slopes that represent the true fitness of each variant (where 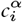 is the y-intercept of this trajectory):

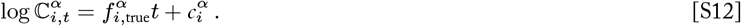

The amount by which the fitness estimate obtained from observed data deviates from the true fitness is given by

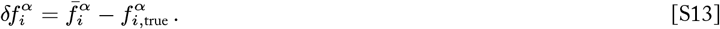

This mis-estimation in fitness is equal to the difference between the ‘rise’ of the observed and true count trajectories over the ‘run’ of the time series (here we only care about the difference in slopes, so we can assume that the two count trajectories are y-shifted to have the same y-intercepts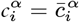 w.l.o.g.):

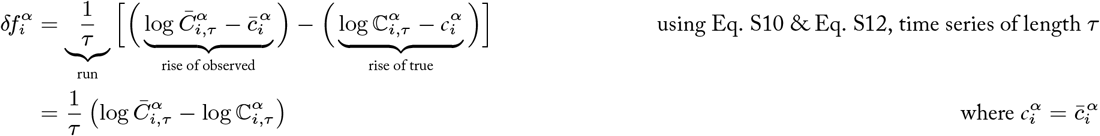

We can evaluate this difference further:

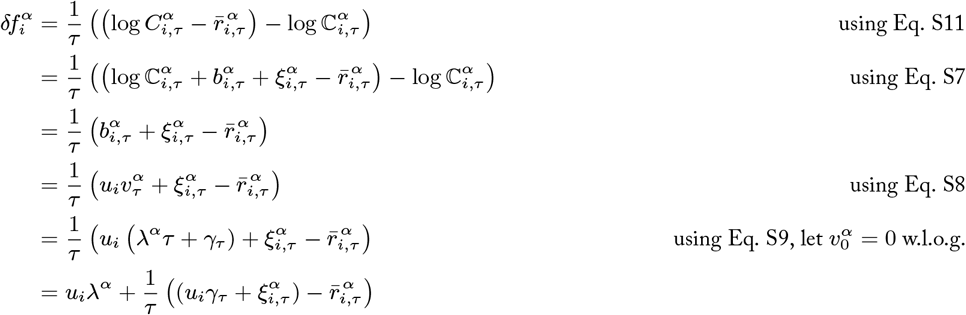

The first term tells us that the error in fitness estimated from observed counts follows from the assay’s trend in bias prevalence, as mediated by the degree of bias susceptibility for the variant in question. The second term is a contribution to this error related to the difference between the observed residual for the final sample and the incidental effects of bias and noise on that sample. This difference can be interpreted as frustration in the linear fit that is *not* accounted for by the effects of bias or noise in barcode processing (i.e., the mapping of true counts to observed counts, Eq. S7), such as deviations from exponential (log-linear) trajectories of abundances in the assay itself. The contribution of the second term is expected to be small and diminishes as the length of the time series increases, such that the error in fitness estimation will be dominated by the first term. That is, the mis-estimation of the fitness of variant *i* is equal (to first order) to the trend in bias prevalence as mediated by that variant’s bias susceptibility:

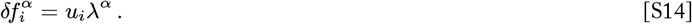

#### S1.4 Bias-corrected counts

We improve the accuracy of fitness estimates by calculating a set of adjusted counts that ‘remove’ the effect of bias from the values that are used to fit a log-linear model. That is, the bias-adjusted count for barcode *i* in sample *t* of assay *α*, denoted by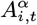, is obtained by subtracting the estimated bias effect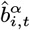 from the observed count for that barcode and sample:

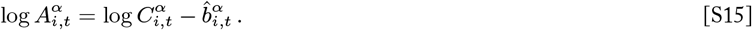

Following our model of bias (Eq. S8), we decompose the estimated bias into estimates for the underlying bias susceptibility and prevalence components:

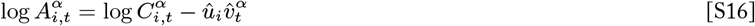

Accurate estimates of the bias components 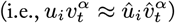 will bring the bias-adjusted counts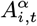 closer to the true counts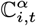 especially when random noise is negligible:

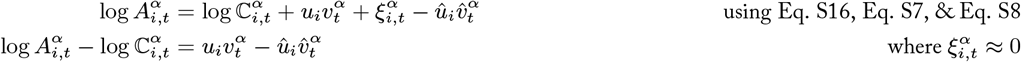

Fitting a linear model to the adjusted counts yields a bias-adjusted estimate of fitness:

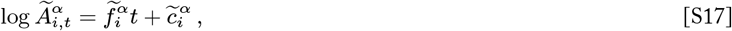

Where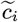 is the y-intercept and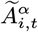 is the predicted adjusted log-count of barcode *i* in sample *t* under the best-fit linear model (the ‘tilde’ notation denotes quantities inferred from the *adjusted* counts data). An adjusted log-count 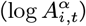 deviates from the corresponding best-fit line by the residual 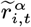:

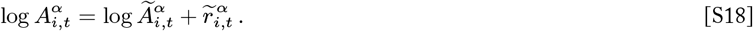

Note that the values of bias-adjusted counts and their corresponding residuals depend on the bias susceptibility and bias prevalence values that are used to compute the adjustment.

**Fig. S1.**
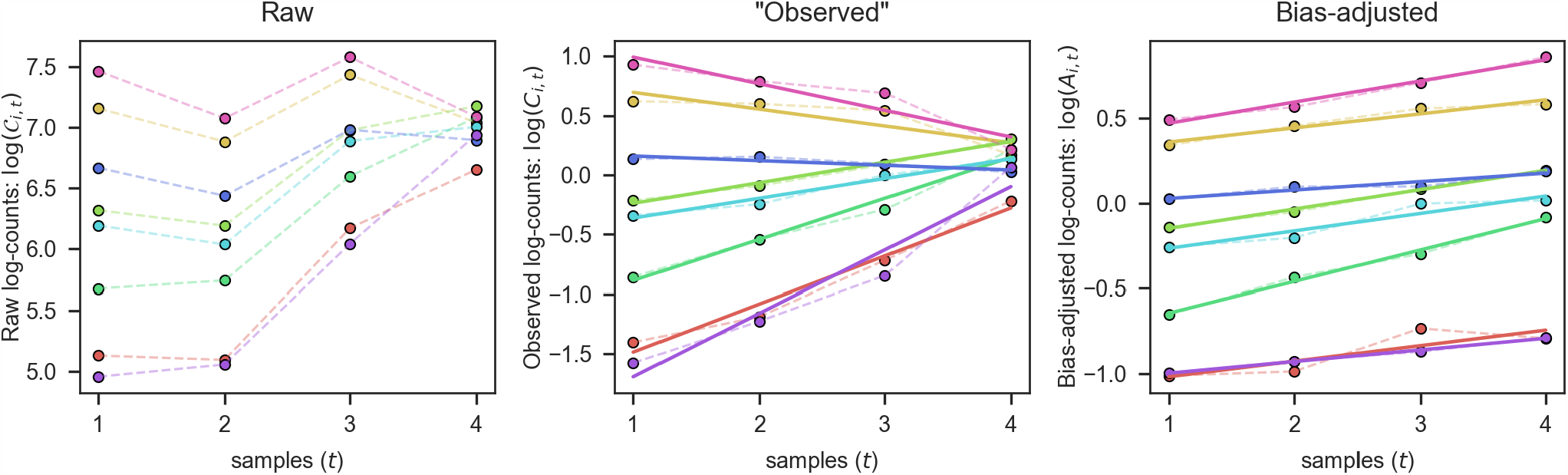
Example of updates to counts data using our method. **(Left)** Raw log-count trajectories (dashed lines) for eight variants (different colors) in the ‘EC2 rep2’ assay from the Kinsler et al. (18) data set are shown (see Supplementary Section S3 for more information). The depicted variants are all members of the near-equal fitness control set used in our case study (see Supplementary Section S3.2), so they are expected to have log-count trajectories with nearly-equal slopes. (Center) Observed log-counts trajectories obtained by normalizing counts to account for varying sequencing depth (dashed lines; Eq. S6) and the corresponding best fit linear regressions (solid lines; Eq. S10) are shown. The depth-normalization makes observed log-count trajectories more linear than the raw trajectories. However, there is large variance in the slopes of the observed log-linear trajectories, which is unexpected for near-equal fitness variants and indicative of systematic bias. (**Right**) Log-count trajectories following bias-adjustment using our method (dashed lines; Eq. S15) and the corresponding best fit linear regressions (solid lines; Eq. S17) are shown. The bias-correction reduces the magnitude of residuals on average and results in fitness estimates that are more accurate, as seen in a decrease in the variance of slopes (i.e., fitness estimates) for this set of near-equal fitness variants.

#### S1.5 Bracket notations

- **Sets:** Curly brackets denote a set of values. A set contains all elements corresponding to the indices written inside the brackets, while the set is associated with a specific element of the index written outside the brackets (if any). For example, 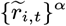 denotes the set of residuals for all variants and all time points associated with a particular assay *α*, whereas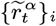 denotes the set of residuals from all assays and all time points for a particular variant *i*.
- **Averages:** Angle brackets denote the arithmetic mean of a set of values, where the ensemble that is averaged includes all elements corresponding to the indices written inside the brackets, and indices written outside the brackets specify elements that are fixed with respect to the average (consistent with the set notation above). For example,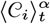 denotes the average count of all variants for a particular sample *t* from a particular assay *α* 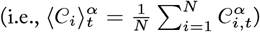.

### S2 Extended Methods: Bias Inference & Correction Algorithm

Our bias inference and correction method proceeds through the following phases:

#### S2.0 Initialization

#### Filtering variants with untrustworthy counts

Variants that have very low or very high abundances in a given assay may have read counts that are impacted by factors other than barcode processing bias, such as high counting error, zero counts in some samples, or deviations from exponential growth (log-linear trajectories). Variant *i* is considered to have ‘*trustworthy*’ data in assay *α* if its average raw count across samples in that assay falls between designated minimum and maximum mean-count thresholds (i.e., 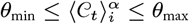 ; threshold values used in our case study are given in Supplementary Section S3.3). To avoid extreme counts unduly influencing the inference of bias components, we exclude data for ‘untrustworthy’ variants from certain parts of our procedure, as described below.

- **Inference of bias prevalence deviations:** Bias prevalence deviation values are inferred on a per-assay basis, using the set of residuals over variants and samples for each assay (i.e., 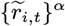 ; see Supplementary Section S2.1b). Variants that are not trustworthy in a particular assay are excluded from the set of residuals used to infer bias prevalence deviations for that assay (although they may be included in other assays where they are trustworthy).
- **Inference of bias susceptibility:** Bias susceptibility values are inferred on a per-variant basis, using the set of residuals from all assays and samples for each variant (i.e., 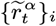 ; see Supplementary Section S2.1a). If a variant does not have trustworthy data in at leastΦassays, then its bias susceptibility is not inferred but rather fixed at zero (the value of Φ used in our case study is given in Supplementary Section S3).
- **Inference of bias prevalence trends:** Bias prevalence trend values are inferred on a per-assay basis, using inferred bias susceptibility values and estimated fitness errors for a pre-designated control set of equal-fitness variants (Supplementary Section S2.2). In order to be included in the trend inference for a particular assay, a control-set variant must be in have trustworthy data for that assay and have met the trustworthiness criteria for bias susceptibility inference (i.e., meeting both criteria outlined above).

##### Assigning weights to data points

We infer bias components and depth-normalization factors using variations of least squares regression, which infers the parameter values that generate adjusted counts with the smallest sum of squared residuals (relative to the best-fit log-linear model; see Supplementary Section S2.1 for details). However, not all counts, and therefore not all residuals, are equally reliable. For example, the counts of low-abundance variants are expected to be more impacted by counting noise (i.e., have a greater variance-to-mean ratio) than those of high-abundance variants. To account for variability in count precision, we use forms of Weighted Least Squares (WLS) regression, in which each residual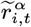 is given weight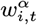 that determines its relative ‘importance’ in the regression.

In WLS regression, it is standard to assign weights equal to the reciprocal of the variance of the observations. Here the relevant observations are log-counts, for which we must determine the variance. Let the raw read count of variant *i* in sample *t* of assay *α* be a Poisson-distributed random variable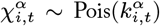. The mean and variance of this count distribution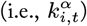 depends on the culture abundance of this variant in this sample, but we assume we do not have this kind of distributional information. Instead, we receive a raw count measurement 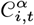. Based on a single measurement, the best estimate of the mean and variance of 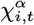 is 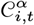 itself. However, our regressions take place in log-count space, so we still need to determine Var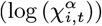. Finding the variance of a non-linear transformation *g*(*X*) of a random variable *X* is non-trivial, but the delta method provides an approximation when the variance of *X* is known:

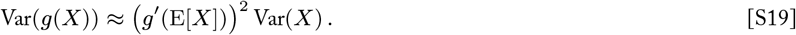

In our case, we have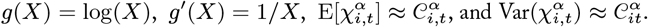 Plugging these into the delta method equation above, we have:

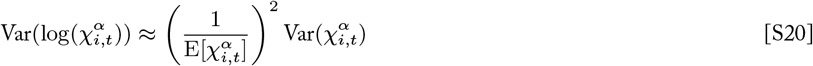

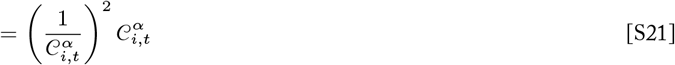

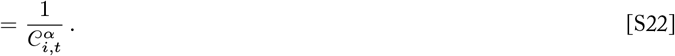

Therefore, the variance of log-count of variant *i* in sample *t* of assay *α* is well-approximated by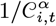.

Defining the weight of a data point as the reciprocal of the corresponding observation’s variance, we have:

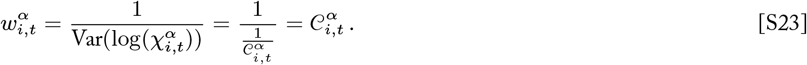

We modify this weight assignment slightly, capping the maximum weight at a designated maximum weight threshold, *W*max:

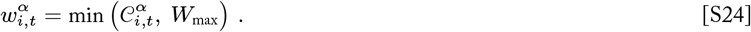

This reflects an assumption that all sufficiently large count observations are equally reliable for our inference purposes.

##### Initializing bias components

Our method infers sets of depth-normalization, bias susceptibility, and bias prevalence values using non-linear least squares optimization, as described in the following sections. Initial values for each of these sets must be specified to serve as initial conditions to the optimization algorithm. Our package allows the user to provide initial values for these sets, but default initializations are presented here:

- **Initial depth-normalization factors:** By default, initial normalization factor values are set to the mean raw counts for the respective samples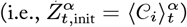. *Note: Depth-normalization factors must not be zero*.
- **Initial bias susceptibilities:** By default, initial bias susceptibility values are drawn from a normal distribution with mean zero and small variance: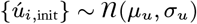 (values used in our case study are given in Supplementary Section S3.3).
- **Initial bias prevalences:** By default, initial bias prevalence values are drawn from a normal distribution with mean zero and small variance: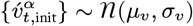 (values used in our case study are given in Supplementary Section S3.3).

*Note: Bias susceptibilities and bias prevalences should not be initialized to zero, as this can cause the gradient of the bias effect* 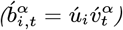 *to be zero, which impairs convergence of the optimization*.

#### S2.1 Inference Stage 1: Inferring bias susceptibilities and bias prevalence ‘deviations’

This stage of the method infers the depth-normalization factors, variant bias susceptibilities, and bias prevalence deviation terms that jointly optimize the log-linearity (i.e., minimize the set residuals) of the correspondingly-adjusted count trajectories. Recall that the estimated effect of bias on an observed count value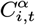 is modeled by the product of bias suceptibility and bias prevalence components (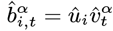, Eq. S8). Subtracting the estimated bias effects from all observed counts yields a set of bias-adjusted counts (Eq. S16), and fitting a linear model to the adjusted counts yields a set of “bias-adjusted residuals”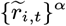 (Eq. S17, Eq. S18). As such, the bias-adjusted residual values that are obtained depend on the bias component values that are used (as well as the normalization factors used to produce the observed counts):

**Fig. S2.**
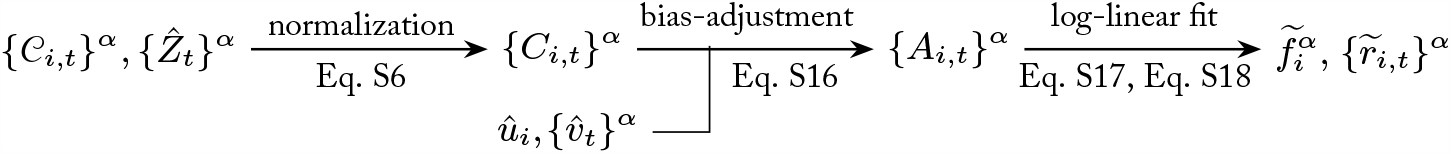
Dependence of bias-adjusted residuals on bias components and depth-normalization factors.

At this stage, we iterate over the following optimization steps (described below) until suitable convergence of the inferred values is reached (as determined by a numerical precision threshold or designated number of iterations).

- **Stage 1a**: Inferring bias susceptibilities
- **Stage 1b**: Inferring bias prevalence deviations
- *Bias component re-normalization step*

#### S2.1a Inferring variant bias susceptibilities

For each variant (with adequate trustworthy data; see Supplementary Section S2.0.1), we infer the bias susceptibility value that minimizes that variant’s consequent bias-adjusted residuals across all samples and assays. In particular, we use numerical non-linear least squares optimization to solve for the susceptibility value that satisfies:

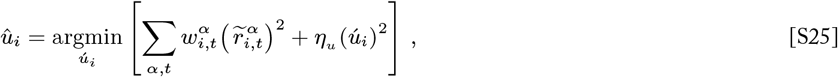

where all bias prevalences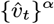 and normalization factors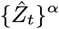 are fixed at their present values. This optimization is a weighted least squares regression with a ridge-like penalty *η*_*u*_ on the magnitude of the susceptibility value. Weighting the regression gives counts with low estimated variance (i.e., more reliable counts) more influence over the susceptibility inference (for more information about weights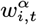 see Supplementary Section S2.0.2). The penalty term favors small susceptibility values and limits outliers.

#### S2.1b Inferring sample bias prevalence deviations

For each assay, we infer a set of per-sample bias prevalence values that minimize the consequent collection of residuals over all time points and variants (with trustworthy data for the given assay; see Supplementary Section S2.0.1). In particular, we use numerical non-linear least squares optimization to solve for the bias prevalence values that satisfy:

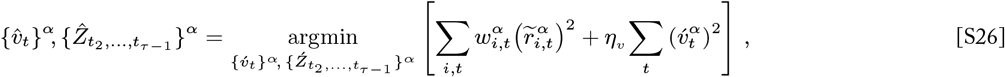

where all bias susceptibilities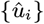 are fixed at their present values. This optimization is a weighted least squares regression with a ridge-like penalty *η*_*v*_ on the magnitude of the prevalence values. Weighting the regression gives counts with low estimated variance (i.e., more reliable counts) more influence over the inference (for more information about weights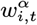 see Supplementary Section S2.0.2).

In this step, the penalty term plays two roles. First, it regularizes the inference, which favors small prevalence values and limits over-fitting. Second, it is important to note that the log-count residuals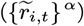 reflect the bias prevalence deviations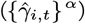 from an underlying linear trend in bias prevalence 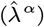 (Eq. S9). As a result, an infinite number of different bias prevalence time series with different slopes but the same deviations will result in the same residuals. This ‘zero mode’ (symmetry) must be resolved in order for the optimization to converge and for the resulting bias prevalence values to not include arbitrary trends. The penalty term resolves this by favoring small bias prevalence values that in turn bias the inference to sequences of prevalences with near-zero slope 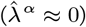.

In this stage of the method, we implicitly enforce that there are no temporal trends in bias prevalence, and therefore inferring bias prevalence values is equivalent to inferring the bias prevalence *deviations* from the near-zero trend 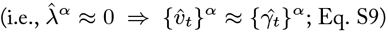. In Stage 2 of the method, we relax this assumption and infer the actual trend in bias prevalence for each assay, using the inferred trend and deviation values to update the absolute bias prevalence estimates for all samples.

Notice that this step optimizes depth-normalization factors jointly with bias prevalence inference. More precisely, we fix the normalization factors for the first and last time points of the assay (*t*_1_ and *t*_*τ*_, respectivley), and we optimize the remaining intermediate time points (*t*_2_, …, *t*_*τ −*1_) along with the bias prevalence values. Fixing the first and last time points prevents the sequence of normalization factors from drifting arbitrarily and impairing convergence due to the interaction of normalization factors and bias prevalence values in determining adjusted counts and consequent residuals.

##### Bias component re-normalization step

We model the effect of bias on a given log-count as the product of bias susceptibility and bias prevalence components (Eq. S8). However, this product is only defined up to a common factor *a*:

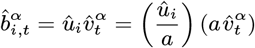

That is, an infininte number of combinations of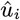,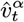, and *a* values can give the same bias effect value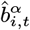. This must be resolved in order for our iterative inference process to converge. We do so by fixing the factor *a* to a particular value and re-normalizing bias component values accordingly during each iteration of Inference Stage 1. In particular, we do the following:

1. Rescale the current bias prevalences: Multiply all bias prevalence values by a common factor *a*, which is defined such that this multiplication rescales the collection of all bias prevalences to have a fixed target standard deviation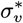 :

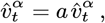

where

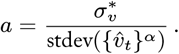
2. Re-normalize the current bias susceptibilities: Divide all bias susceptibility values by the common factor *a* defined above, which renormalizes the susceptibility values to the newly rescaled prevalence values and maintains the absolute bias effects:

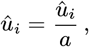

#### S2.2 Inference Stage 2: Inferring bias prevalence trends

Coming out of Stage 1, we have obtained a set of inferred bias susceptibility value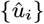 for all variants and sets of bias prevalence values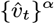 for all assays. In Stage 1, our regularization enforced that inferred bias prevalence values have negligible temporal trends (i.e., near-zero slopes) within each assay, so these values are better interpreted as sets of approximate bias prevalence deviation values 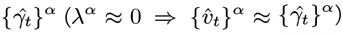. In Stage 2, we infer the actual trend in bias prevalence for each assay, using the inferred trend and deviation values to determine the absolute bias prevalence for all samples.

To infer bias prevalence trends, we make use of the relationship between fitness misestimates (before bias correction) and bias susceptibility (Eq. S14):

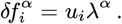

This relationship tells us that the trend in bias prevalence for an assay can be estimated by regressing misestimates of fitness from that assay against bias susceptibilities for a set of variants for which these values are known.

In general, fitness misestimate information is not available in the context of fitness assays. However, our method requires that the assays under consideration include a designated control set of equal-fitness variants, which we denote as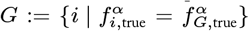. That is, the variants in the control set *G* are assumed to share the same ‘true’ fitness,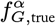 The true fitness of the control variants can be estimated by taking the average of the observed fitness estimates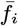 among the control variants:

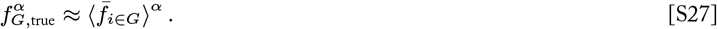

Then the misestimate of fitness for each variant in the control set can be estimated by

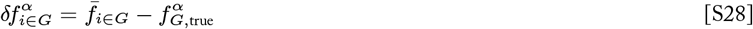

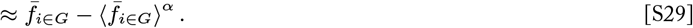

Therefore, inclusion of a control set of equal-fitness variants avails a set of variants for which both fitness misestimates and bias susceptibilities can be obtained.

We use these values to estimate the trend in bias prevalence for each assay using ordinary least squares regression:

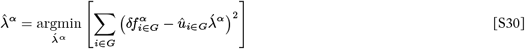

At this point, we have inferred sets of bias prevalence deviation values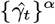(from Stage 1) as well as bias prevlance trend values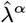 (from Stage 2) for each assay. These temporal trends in bias were not resolvable from the residuals-based inference in Stage 1, but we can now compute bias prevalence values that incorporate this temporal information and reflect the actual effects of bias on each assay (Eq. S9):

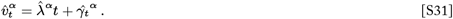

Since it is the slope of bias prevalences not their absolute value that ultimately impacts fitness estimates, we conclude this stage by centering the bias prevalence values for each assay (such that the y-intercept is near zero) without loss of generality:

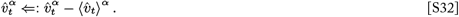

#### S2.3 Outputting bias-corrected counts and fitness estimates

At the end of the iterative inference stage, we possess a set of inferred bias susceptibility values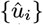 for all variants, sets of (absolute) bias prevalence values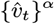 for all assays, and inferred depth-normalization factors for all samples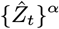. We use the final inferred values to compute a final set of bias-adjusted counts for all variants and all assays (Eq. S16). Log-linear models fit to these final bias-adjusted counts yield final bias-adjusted fitness estimates (as well as final bias-adjusted residuals; Eq. S17). The output of our algorithm includes the final bias-corrected counts and fitness estimates, as well as other metadata.

### S3 Case Study

#### S3.1 Data set

We applied our bias inference and correction method to the yeast bulk fitness assay data from Kinsler et al. (18). These fitness assays were performed with barcoded mutants (variants) isolated from a previous evolution experiment (16), where were competed against a constructed reference strain with a restriction site in the barcode region (17). This library was assayed in a total of 45 environments (i.e., abiotic culture conditions). These environments include instances of the limited glucose “evolution condition” (EC) in which the library’s variants evolved, as well as a range of environments that each feature a small perturbation to this condition, such as subtle changes in the amount of glucose, carbon source, or stressors present (e.g., salt). Refer to Kinsler et al. (18) for additional information about yeast strains, variant mutations, and assay culture conditions.

In this case study, we restrict our analysis to data from environments for which all corresponding samples have a total sequencing depth of at least 100,000 reads. We analyze data from 35 assays across 15 environmental conditions, as detailed in the table below. For concision, the main text only displays results for assays from the EC environments, but data from all 35 assays and all 15 environments were used for inference. Results for all environments can be found in Supplementary Section S3.4.

**Fig. S3.**
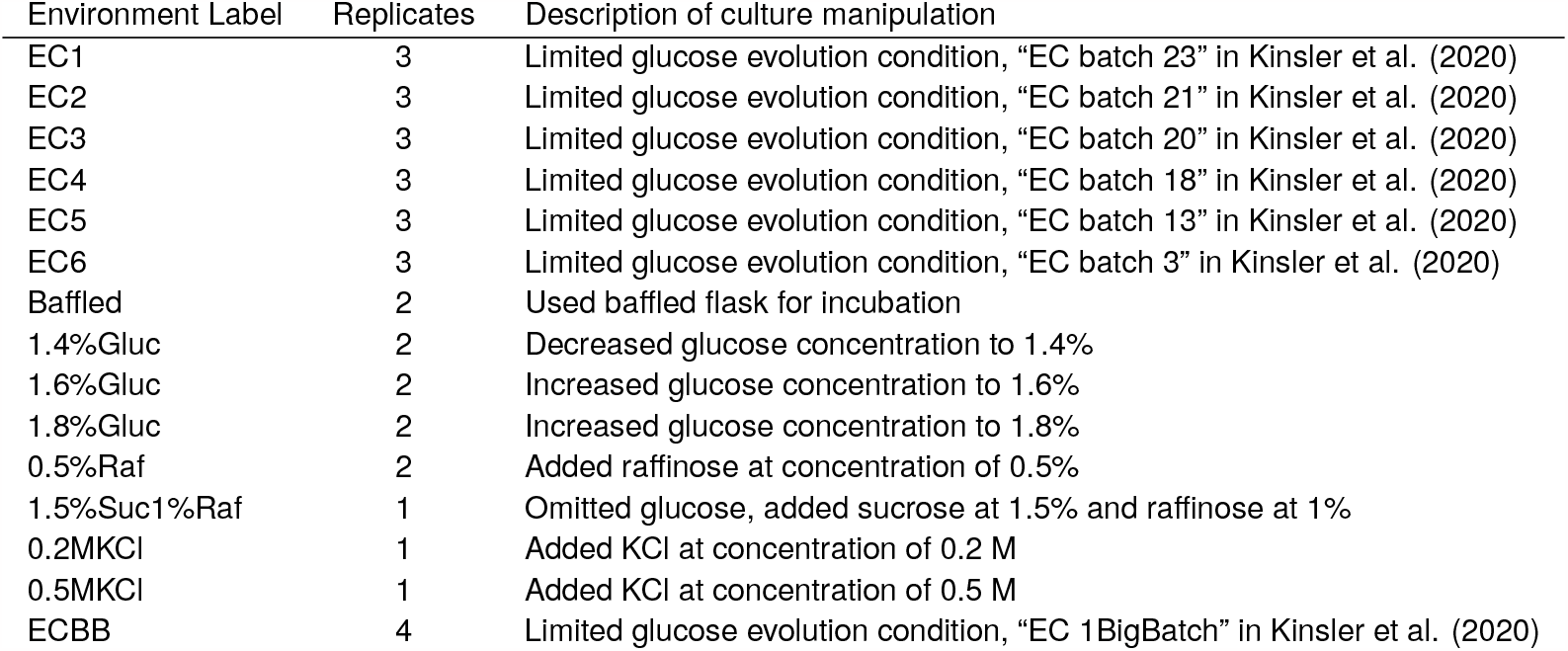
Assay conditions included in this analysis.

#### S3.2 Control set of equal-fitness variants

Our method requires that a subset of variants with near-equal fitnesses be designated for use as a control set (Supplementary Section S2.2). Ideally, the variant library would be designed with a multiply-barcoded but otherwise identical set of strains for this purpose. Here, however, we are applying our bias correction method to fitness assays that were conducted without this consideration in mind, so we must identify a suitable subset of variants with near-equal fitnessses *a posteriori*.

The Kinsler et al. (18) data set includes 549 variants, each of which represents a mutation that arose in response to the limited glucose evolution conditions. A large number of these mutations fall in a small number of genes, including *GPB2* and *PDE2*. In addition, many mutants are diploids resulting from whole-genome duplication events but have otherwise very similar genetic backgrounds. These classes of shared mutation targets offer candidate groups for variants with near-equal fitness effects.

To select groups of putatively near-equal fitness variants, we carried out the following outlier exclusion procedure for each of the aforementioned mutation classes (i.e., *GPB2, PDE2*, and diploids), First, we computed the fitness of each variant using the uncorrected observed counts for each assay (Eq. S10), obtaining a vector of 35 fitness estimates for each variant. We also computed the vector of mean fitness estimates for each mutation class. We computed the Mahalanobis distance of each variant’s fitness vector from its mutation class’s mean vector, and variants with distances beyond the 90th quantile of the associated chi-squared distribution were called as outliers and excluded. The remaining variants constituted a group of putatively near-equal fitness variants for the given mutation class.

To gauge the validity of these putative groups, we computed the SVD spectrum of the matrix of fitness mis-estimates (Eq. S13; variants as rows, assays as columns) for each mutation class (Figure S4). If a group of variants has the same true fitness, then differences in observed fitness estimates among the group are only attributable to systematic bias (assuming the effects of random noise on fitness estimates are small), and the effects of a single bias mode (consistent with our model of bias) result in matrix of fitness *mis*-estimates of rank one. Therefore, a matrix of fitness *mis*-estimates with a SVD spectrum that is dominated by the first mode is consistent with the corresponding group of variants having nearly equal true fitnesses. We found that the first mode accounted for at least 60% of the variance in fitness mis-estimates for the diploid, *GP2*, and *PDE2* groups (Figure S4), which supports the suitability of these variants groups as near-equal fitness sets. In our case study, we used the largest of these groups—the diploids—as the control set for our method, and we considered the *GP2* and *PDE2* groups as validation groups (see Results, Figure 4).

**Fig. S4.**
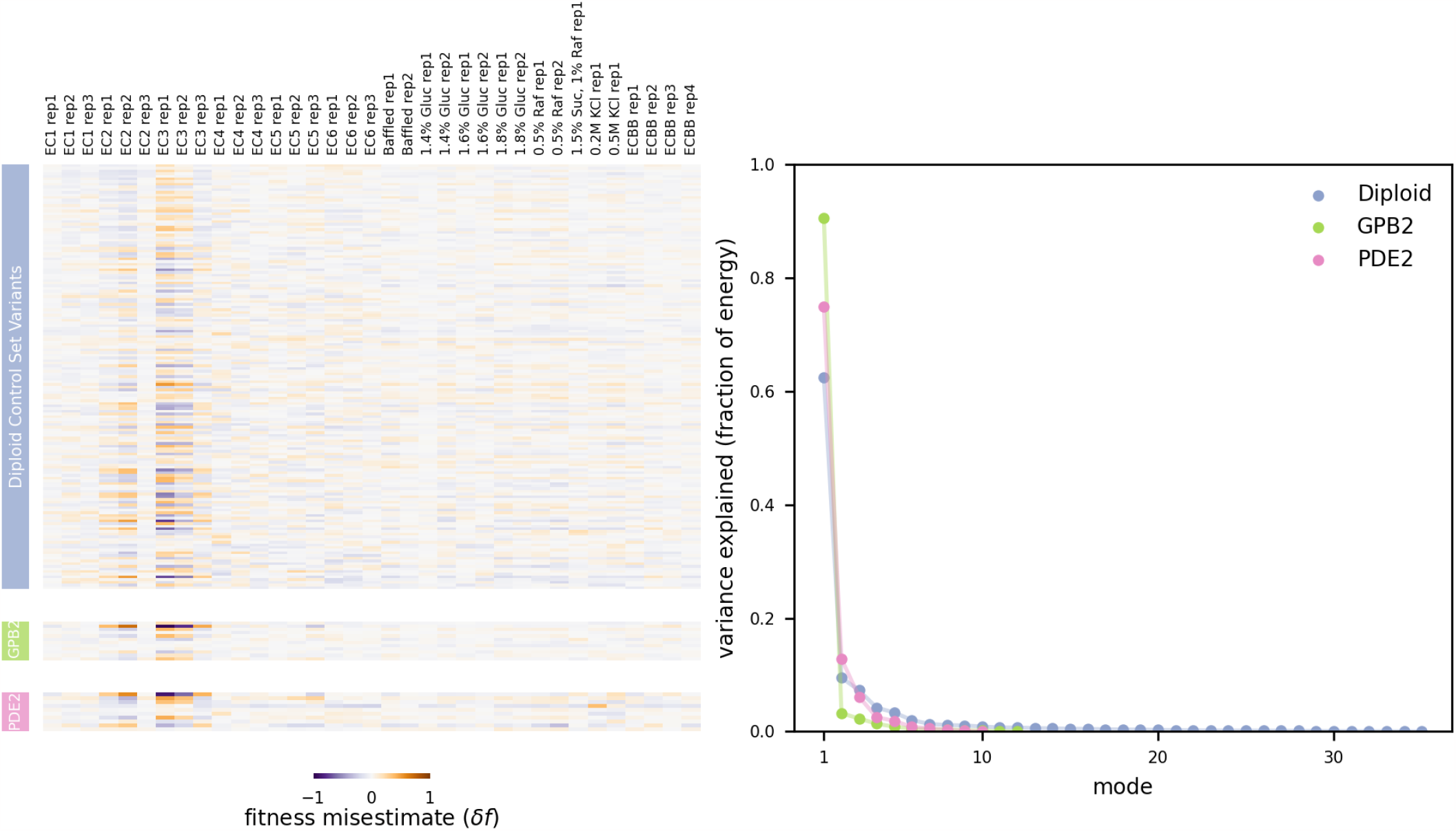
**(Left)** The heatmaps depict the matrix of fitness misestimates for each variant (row) in each assay (column) for the diploid (top, blue), GPB2 (middle, green), and PDE2 (bottom, pink) mutation classes. (**Right**) The SVD spectrum for each of the fitness misestimate matrices (shown at left) is plotted. The spectrum for each mutation class is dominated by the first mode, which is consistent with a single bias mode accounting for the observed fitness misestimates.

#### S3.3 Method parameterization

In this case study, we executed our method using the following parameters:

**Fig. S5.**
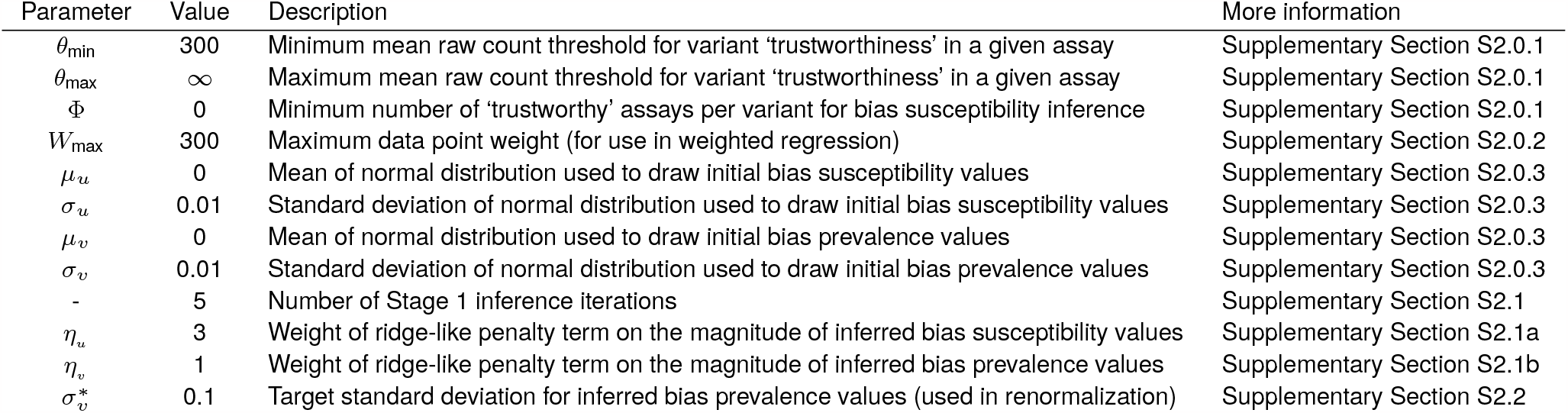
Parameterization of our method for the case study analysis.

#### S3.4 Supplementary Results

**Fig. S6.**
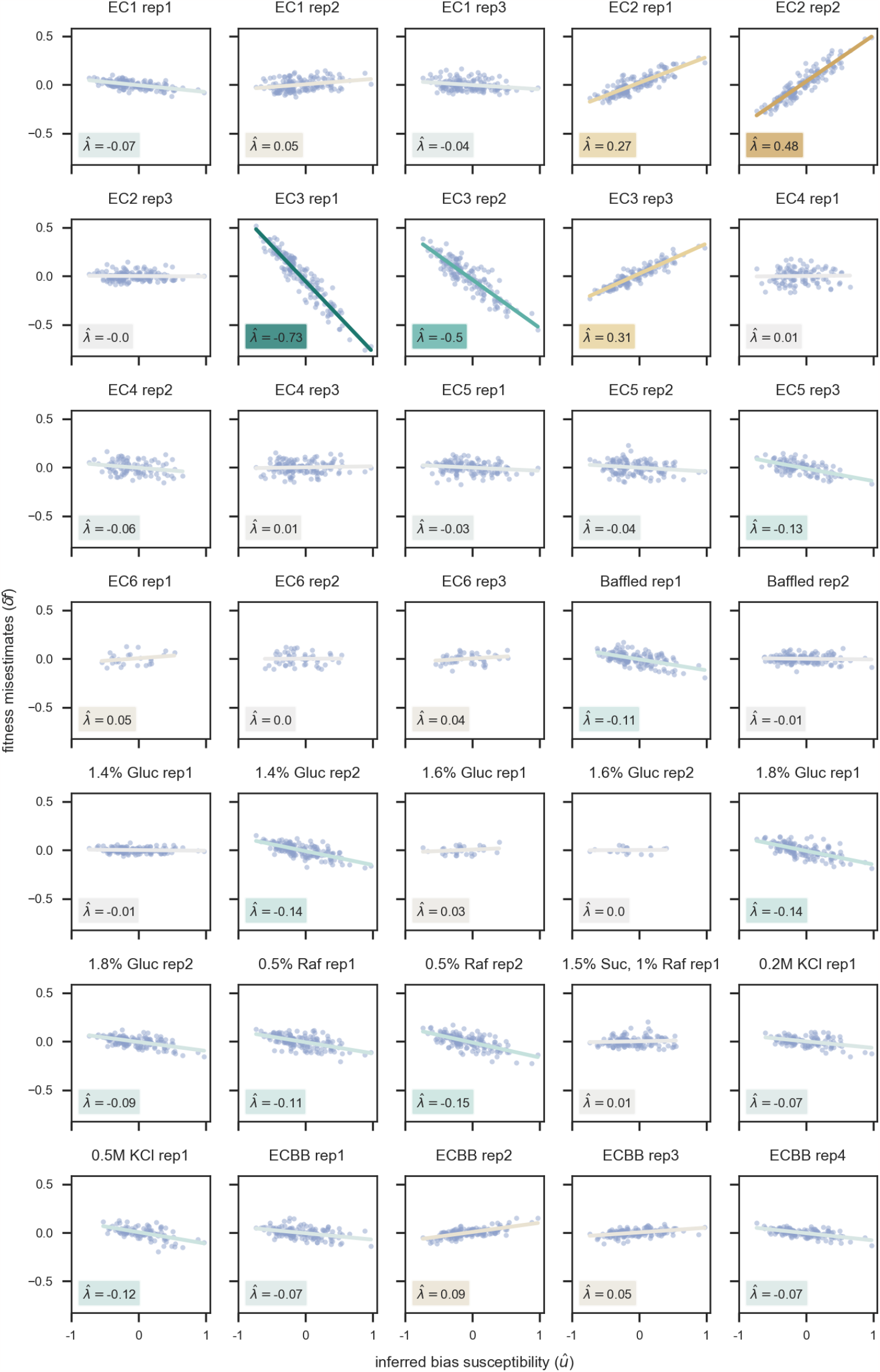
Inference of bias prevalence trends (Algorithm Stage 2). The trend in bias prevalence for an assay can be estimated by regressing misestimates of fitness from that assay against bias susceptibilities for a control set of variants for which these values are known (Eq. 3, Supplementary Section S2.2). Each panel presents the results of this regression for one of the 35 assays in the Kinsler et al. (18) case study data set. Each point represents a variant in the Diploid control set (Supplementary Section S3.2). Only control variants that are ‘trustworthy’ with respect to bias susceptibility inference (Supplementary Section S2.0.1) in a given assay are included in that assay’s regression. The best fit line for each assay is shown, the slope of which corresponds to the inferred bias prevalence trend value 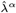 for the assay (values listed in the bottom left of each panel).

**Fig. S7.**
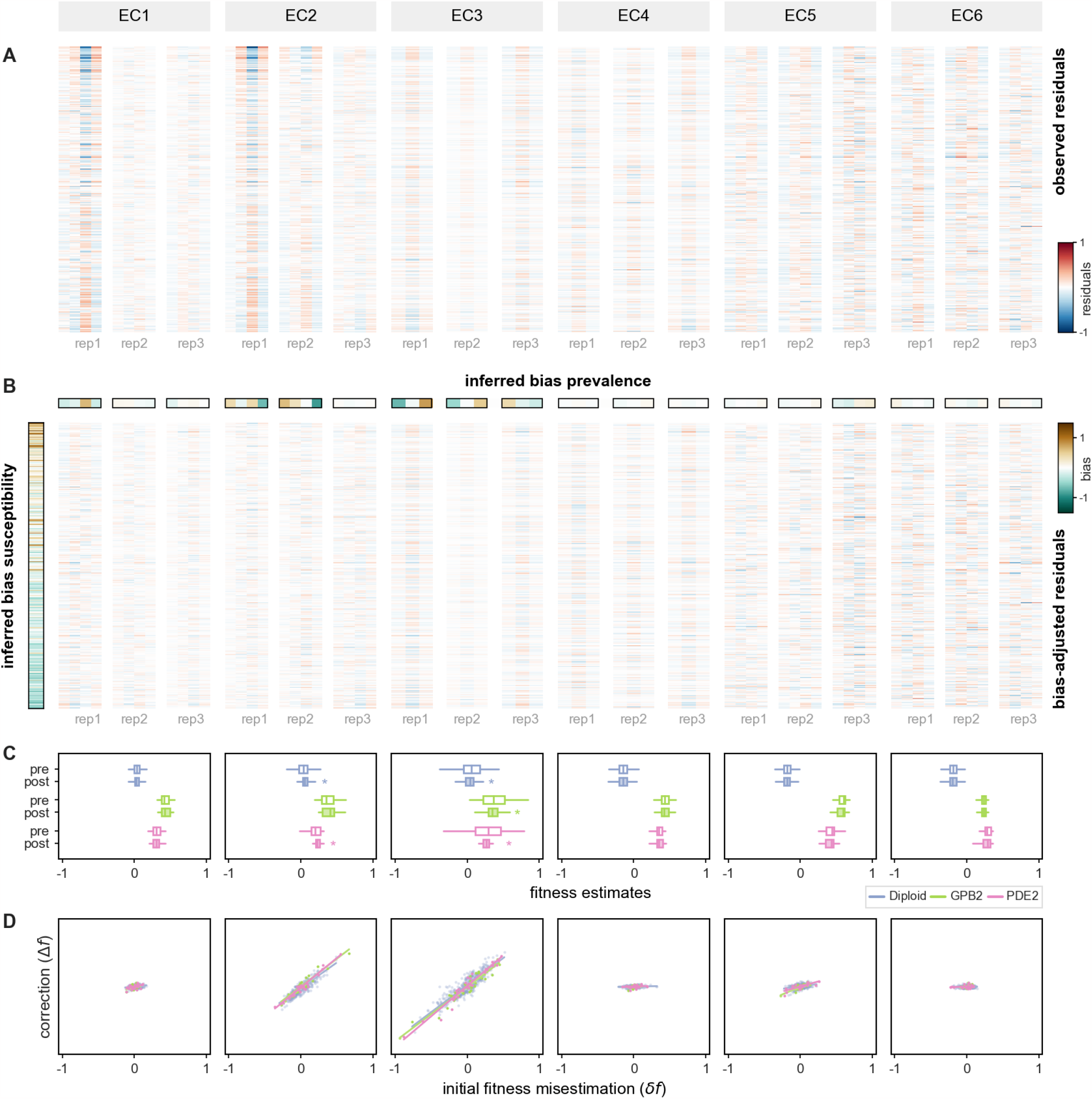
Residuals and fitness estimates before and after bias correction (Part 1/3). We applied our method to the counts data from Kinsler et al. (18) (Supplementary Section S3). This figure presents data and results for 18 assays across six environmental conditions (labeled by the gray boxes, top). These are the same data and results shown in Figure 2 and Figure 4 in the main text, presented side-by-side here for easier comparison of residuals before and after bias-correction. (**A**) The heatmap shows the residuals of linear fits to observed log-count trajectories before bias-correction. (**B**) The bias susceptibility and bias prevalence values inferred by our method are shown in the outlined bands to the left of and above the table, respectively. The post-correction residuals of log-linear fits to the corresponding bias-corrected counts are shown in the heatmap. (**C**) The pre- and post-correction distributions of fitness estimates (white and shaded boxplots, respectively) are shown for each reference group of variants across six growth conditions. Statistically significant decreases in the variance of fitness estimates (as determined by Levene’s test for equal variances; *p* < 0.05) are denoted by *. (**D**) The correspondence between each variant’s initial fitness misestimation using counts before bias correction 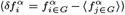 and the change in its fitness estimate following bias-correction 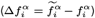is depicted using scatter plots for each growth condition (each point represents a variant).

**Fig. S8.**
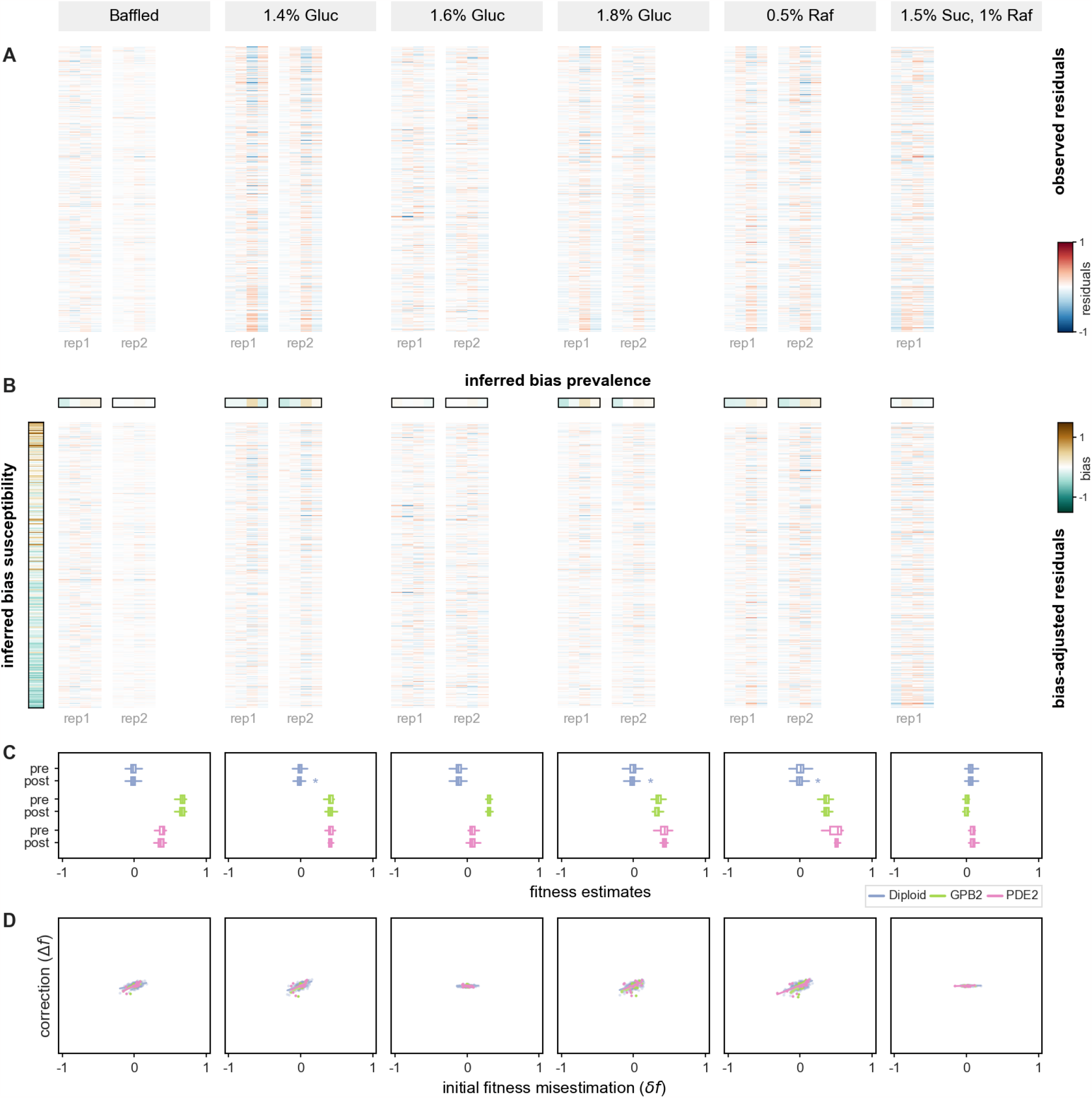
Residuals and fitness estimates before and after bias correction (Part 2/3). We applied our method to the counts data from Kinsler et al. (18) (Supplementary Section S3). This figure presents data and results for an additional 11 assays across six environmental conditions (labeled by the gray boxes, top) that were not depicted in the main text. (**A**) The heatmap shows the residuals of linear fits to observed log-count trajectories before bias-correction. (**B**) The bias susceptibility and bias prevalence values inferred by our method are shown in the outlined bands to the left of and above the table, respectively. The post-correction residuals of log-linear fits to the corresponding bias-corrected counts are shown in the heatmap. (**C**) The pre- and post-correction distributions of fitness estimates (white and shaded boxplots, respectively) are shown for each reference group of variants across six growth conditions. Statistically significant decreases in the variance of fitness estimates (as determined by Levene’s test for equal variances; *p* < 0.05) are denoted by *. (**D**) The correspondence between each variant’s initial fitness misestimation using counts before bias correction 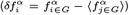 and the change in its fitness estimate following bias-correction 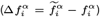 is depicted using scatter plots for each growth condition (each point represents a variant).

**Fig. S9.**
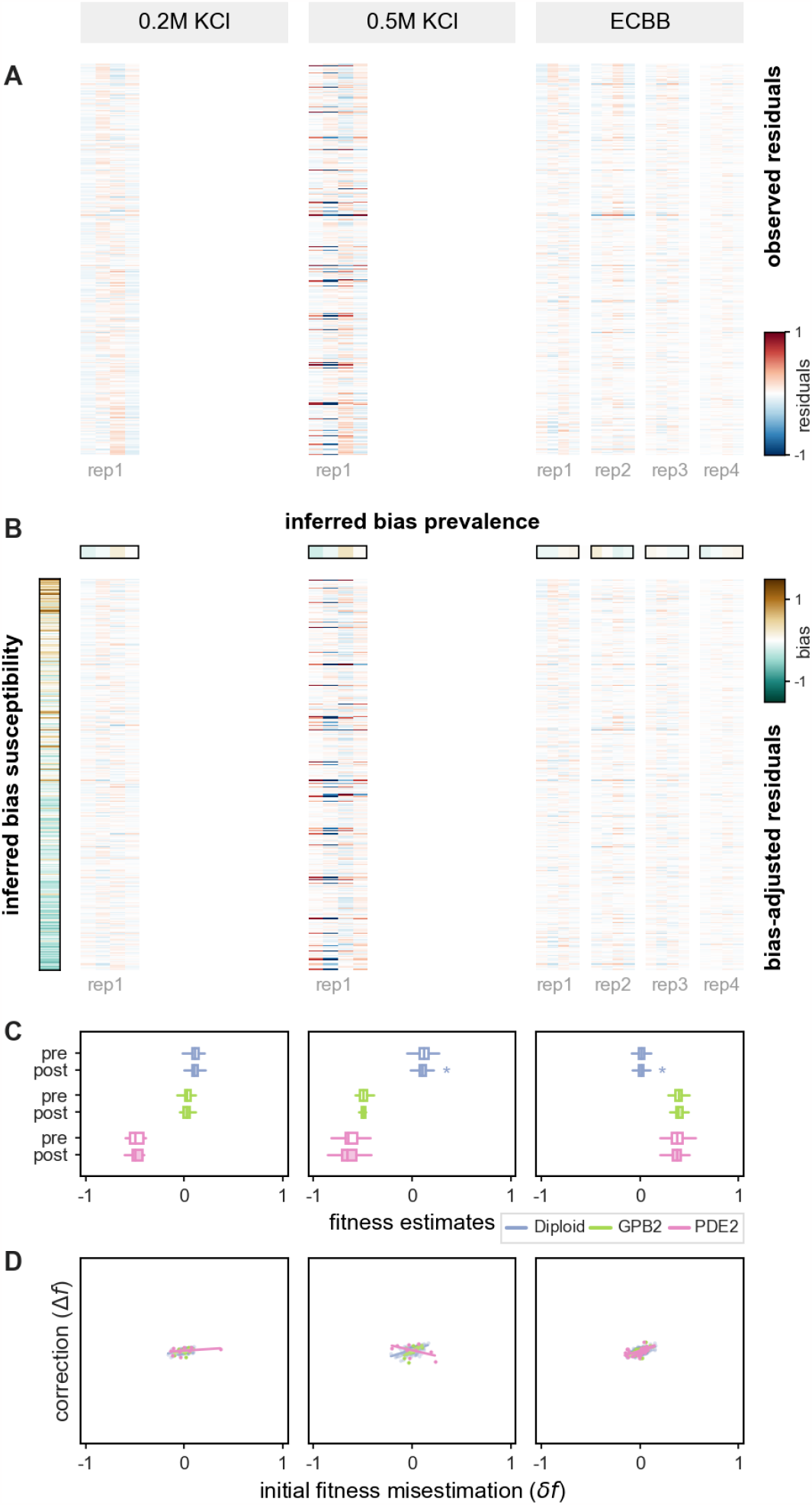
Residuals and fitness estimates before and after bias correction (Part 3/3). We applied our method to the counts data from Kinsler et al. (18) (Supplementary Section S3). This figure presents data and results for an additional six assays across three environmental conditions (labeled by the gray boxes, top) that were not depicted in the main text. (A) The heatmap shows the residuals of linear fits to observed log-count trajectories before bias-correction. (**B**) The bias susceptibility and bias prevalence values inferred by our method are shown in the outlined bands to the left of and above the table, respectively. The post-correction residuals of log-linear fits to the corresponding bias-corrected counts are shown in the heatmap. (**C**) The pre- and post-correction distributions of fitness estimates (white and shaded boxplots, respectively) are shown for each reference group of variants across six growth conditions. Statistically significant decreases in the variance of fitness estimates (as determined by Levene’s test for equal variances; *p* < 0.05) are denoted by *. (**D**) The correspondence between each variant’s initial fitness misestimation using counts before bias correction 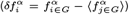 and the change in its fitness estimate following bias-correction 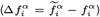 is depicted using scatter plots for each growth condition (each point represents a variant).

